# Multiple Sclerosis Drug Fingolimod Exhibits Antibacterial Activity through Bacterial Membrane Permeabilization

**DOI:** 10.64898/2026.03.02.709040

**Authors:** Antara Syam, Benjamin Rees, Sebastian Cuervo, Fengtian Xue, Alexander Sodt, Ekaterina M Nestorovich, Tatiana K Rostovtseva, Sergey M Bezrukov, John S Choy

## Abstract

Although receptor-mediated mechanisms account for the therapeutic action of numerous FDA-approved drugs, emerging evidence suggests that many of these therapeutics have off-target antimicrobial activities. One example is fingolimod, an immunomodulator used to treat multiple sclerosis, that has been reported to have antimicrobial effects associated with membrane permeabilization. Yet the molecular mechanism by which fingolimod alters bacterial membranes remains unknown. As a cationic amphiphilic drug (CAD), fingolimod is comprised of both hydrophobic and positively charged regions that can enable membrane interactions. We show that fingolimod compromises membrane integrity in *E. coli* and *P. aeruginosa*, contributing to its antimicrobial activity. To determine how fingolimod disrupts membrane integrity, we used planar lipid bilayer electrophysiology with phospholipid compositions mimicking *E. coli* membranes. Using gramicidin A channels as molecular biosensors, we show that fingolimod alters both mechanical properties and surface charge of lipid bilayers at concentrations that have antimicrobial effects. At higher concentrations, fingolimod directly permeabilizes lipid bilayers, as revealed by conductance measurements and Bilayer Overtone Analysis. Molecular dynamics simulations correlate fingolimod’s preference for pore-favoring curvature with its strong interactions with lipids and trans-leaflet translocation. These findings establish a molecular mechanism for fingolimod’s off-target activity and provide a starting point for understanding how some CAD structures can drive membrane-specific effects that compromise bacterial physiology.

**Importance:** Many commonly prescribed drugs, beyond their primary action via receptor targets, modify cell membranes. A mechanistic understanding of how these drugs interact with bacterial membranes will have a significant impact on drug design and on the evaluation of potential side effects. Furthermore, the emerging need for new antimicrobial drugs has led to increased interest in drug repurposing. Elucidating the molecular mechanisms of these compounds’ interactions with bacterial membranes can ultimately provide critical insights into redesigning existing drugs as antimicrobials and into identifying unintended membrane-related effects that may contribute to their therapeutic or off-target effects.

## Introduction

Fingolimod (FTY720), an FDA-approved treatment for relapsing multiple sclerosis, acts as an antagonist of sphingosine-1-phosphate (S1P) receptors, leading to lymphocyte sequestration and immunomodulation (1). Structurally, fingolimod belongs to a class of therapeutics known as cationic amphiphilic drugs (CADs), which have a positively charged amine group and a hydrophobic region. These characteristics enable adequate solubility in body fluids while allowing penetration into lipid membranes (2). Beyond its intended therapeutic mechanism, fingolimod, like other CADs, may have off-target interactions with cellular membranes that remain poorly understood, and some members are known to induce phospholipidosis (2–4). Our previous work, along with studies by other labs, shows that antidepressants, which are also CADs, display antimicrobial effects (5, 6).

Recent studies have revealed unexpected antimicrobial properties of fingolimod, proposed to be associated with perturbation of bacterial quorum-sensing systems and membrane permeabilization (7–10). Furthermore, fingolimod displays synergistic effects with antibiotics such as doripenem and colistin, enhancing their efficacy against resistant bacteria, including carbapenem-resistant *Escherichia coli* (CREC) and *Klebsiella pneumoniae* (11). For example, fingolimod combined with doripenem significantly reduced the expression of resistance-related genes, inhibited bacterial motility, and improved survival in infection models (11). Fingolimod combined with colistin disrupted bacterial membranes, increased oxidative stress, and altered fatty acid metabolism, leading to significant bactericidal effects both in vitro and in vivo (12). Furthermore, fingolimod displays bactericidal activity against gram-positive bacteria like *S. aureus* and *C. perfringens*, where it inhibits biofilm formation and eradicates persister cells (12, 13).

Although fingolimod displays antimicrobial activity and has been shown to disrupt bacterial membranes, the biophysical mechanisms underlying its membrane-permeabilizing effects remain poorly understood. We initially identified fingolimod in a small-scale screen of CADs with bactericidal activities. Using a combination of cellular, biophysical, and computational approaches, we demonstrate that fingolimod alters the properties of planar lipid membranes and induces membrane permeability at concentrations that exhibit antimicrobial activity. Using planar lipid membranes and MD simulations, we show that fingolimod can form distinct membrane pores. These findings provide a molecular mechanism for fingolimod’s antimicrobial activity and suggest potential strategies for developing and repurposing new membrane-targeting antibiotics based on the fingolimod scaffold.

## Results

### A screen of 25 CADs reveals antimicrobial activity of fingolimod

In previous work, it was reported that antidepressants displayed antimicrobial activity (8, 10, 11). Given their characteristic CAD structures, we performed a small-scale screen of primary, secondary, and tertiary amine CADs to assess their effects on *E. coli* growth. We assembled a library of 25 compounds (Table S1), treated *E. coli* with 40 μM of each compound, and measured the growth profile over a 24 h period. We observed a subset of drugs with ∼10% decrease in total growth compared to untreated cells. However, fingolimod was extremely toxic and prevented the growth of cells completely (Fig. 1A). We selected fingolimod for follow-up experiments and tested growth as a function of several concentrations of the compound. Even at lower concentrations (10 μM), a modest reduction of growth was observed (Fig. 1 B, D). To determine whether this growth effect was specific to *E. coli* or more broadly to other bacterial species, we tested effects on the growth of *P. aeruginosa*, another clinically relevant gram-negative bacterium. Similar to the *E. coli*, treatment with increasing concentrations of fingolimod (10 and 40 μM) led to a progressive reduction in *P. aeruginosa* growth (Fig. 1C, D). These findings demonstrate that fingolimod exerts antimicrobial activity in a concentration-dependent manner.

**Figure 1.**
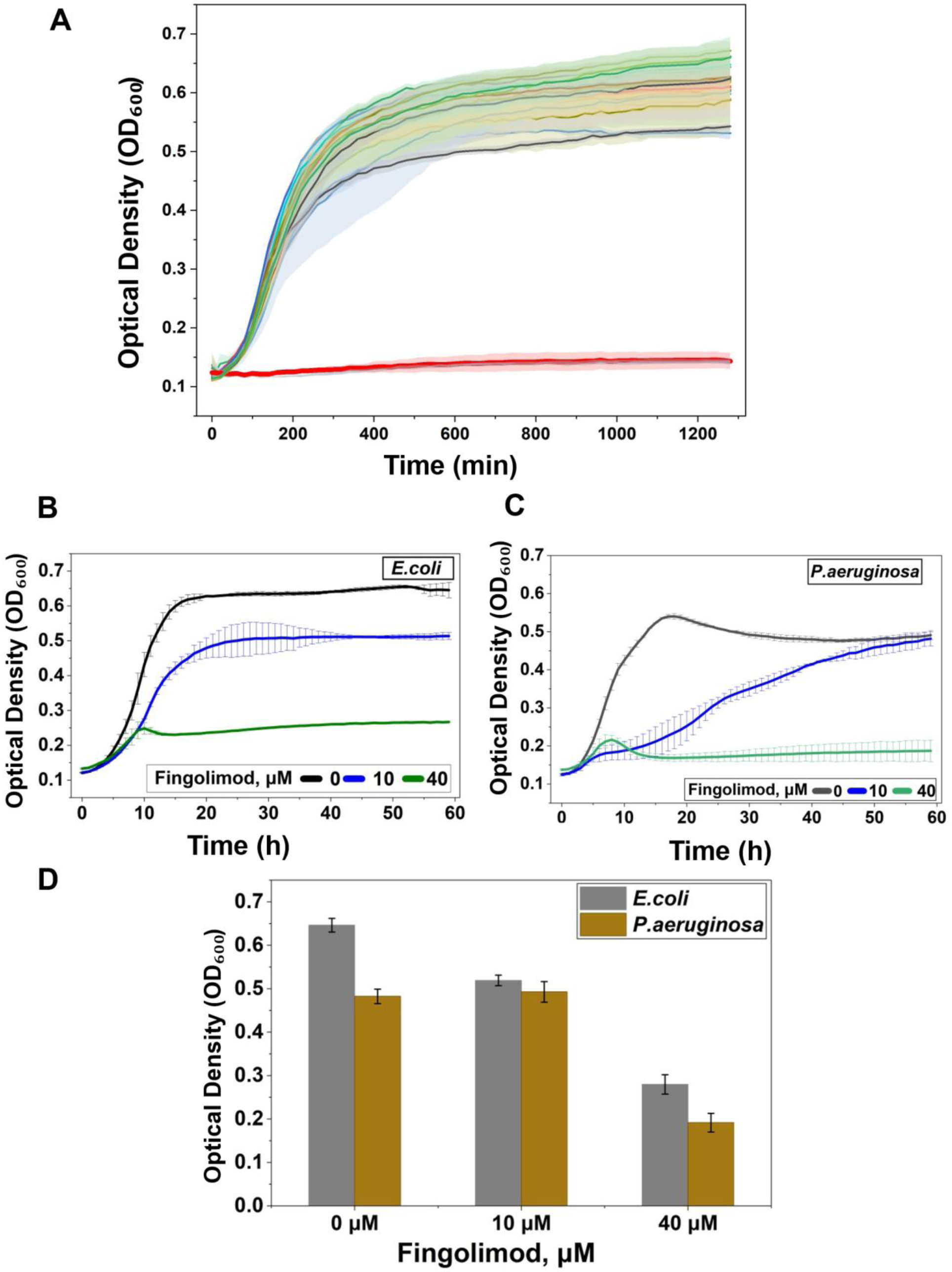
Fingolimod exhibits strong antimicrobial activity and inhibits growth of *E. coli* and P. aeruginosa. **(A**) Growth inhibition screen of 25 cationic amphiphilic drugs (CADs) against *E. coli*. Bacterial cultures were treated with 40 µM of each compound and OD₆₀₀ was measured over a 24-hour period. While many CADs had minimal to moderate effects, fingolimod (FTY720) showed the most potent growth inhibition, nearly abolishing bacterial growth across the time course. **(B–C)** Concentration-dependent inhibition of bacterial growth by fingolimod. *E. coli* (**B**) and *P. aeruginosa* (**C**) were grown in LB medium supplemented with increasing concentrations of fingolimod (0, 10, and 40 µM). OD₆₀₀ was recorded every 10 minutes over 24 hours. Fingolimod inhibited growth in a dose-dependent manner in both bacterial strains, with *E. coli* showing slightly greater sensitivity. **(D)** End-point optical density analysis of bacterial growth after 24 hours in the presence of fingolimod. Bar graphs show mean OD₆₀₀ values ± SD for *E. coli* and *P. aeruginosa* at each concentration tested. A clear dose-dependent reduction in bacterial growth is observed, consistent with the kinetic growth curves.

### Gramicidin A as a molecular biosensor to study CAD membrane interaction

Gramicidin A (grA) serves as an extremely sensitive molecular biosensor of lipid bilayer mechanics (14). GrA is a short linear peptide (15 amino acids) that forms ion channels upon dimerization of its two monomers incorporated into opposing membrane leaflets (15). The dimeric grA channel (∼2.2 nm) (16) is shorter than the typical bilayer thickness (17), resulting in a hydrophobic mismatch (18–20). Therefore, channel formation requires local membrane bending, making grA highly responsive to bilayer mechanical properties (16, 21). The schematic in Fig. 2A illustrates how cationic amphiphiles can alter lipid bilayer properties, thereby modifying grA channel conductance, lifetime, and opening frequency under application of voltage. GrA channel conductance is defined by the density of K^+^ ions at the channel entrance, which, in turn, depends on the membrane surface charge and the bulk electrolyte concentration (22). When amphiphilic fingolimod is added to the planar membrane, it partitions into the bilayer and reduces membrane negative surface charge via its positively charged amine group, repelling the K⁺ ions from the bilayer surface. This reduces the cation concentration near the channel entrance, resulting in decreased channel conductance (22). The lifetime of the grA conducting dimer depends on several membrane parameters, such as bilayer thickness, spontaneous curvature, and lipid packing stress (14, 23, 24). Multiple studies showed that grA channel parameters could report on amphiphilic drug-induced modifications of bilayer electrostatics and hydrophobic core properties (23, 25, 26).

**Figure 2.**
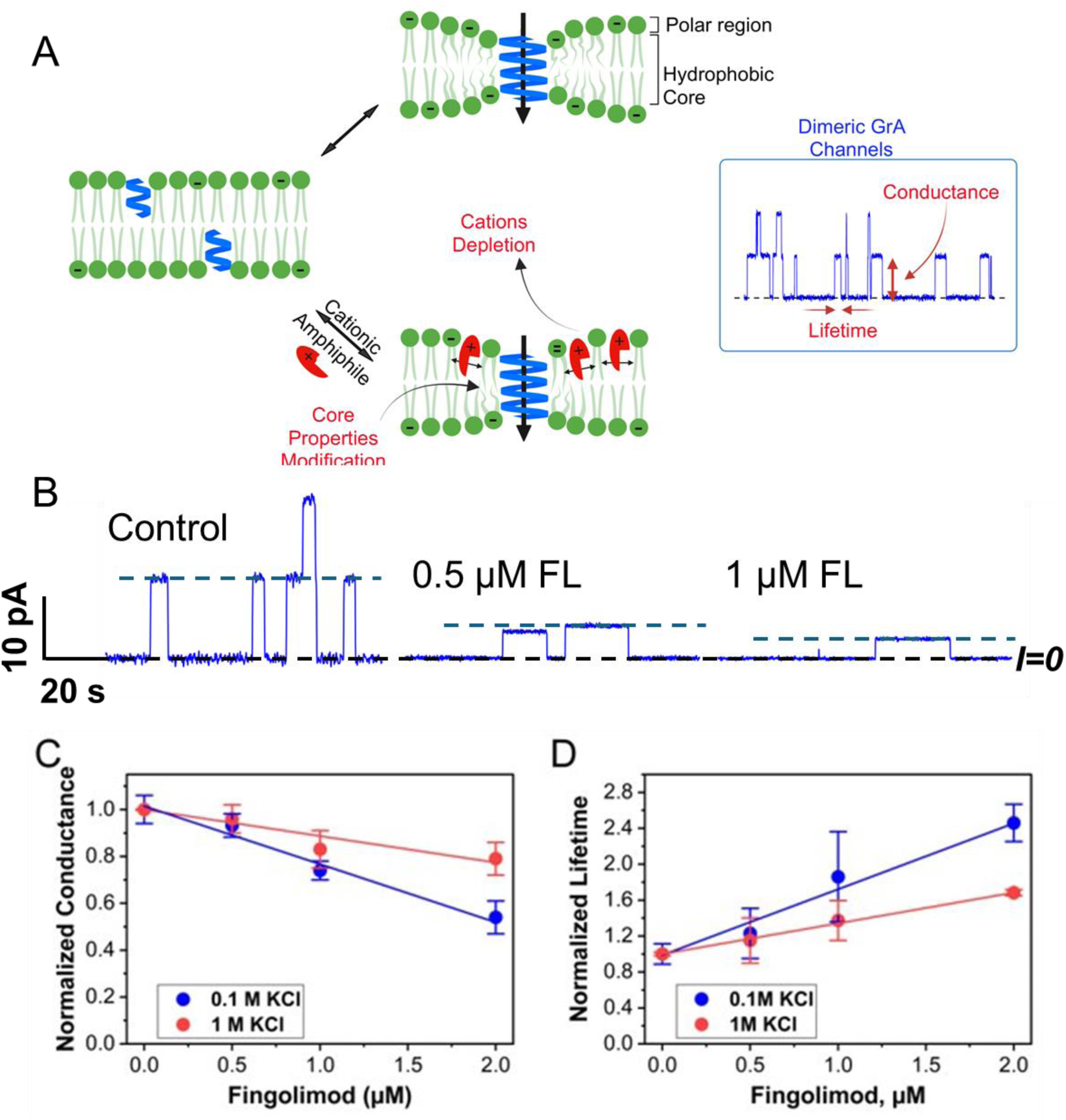
Schematic of GrA channel formation and recording. Fingolimod decreases GrA conductance and increases lifetime. (**A**) GrA is a short linear peptide that forms ion channels upon dimerization of two monomers from opposing lipid bilayer leaflets. Cationic amphiphiles can alter bilayer properties, thereby modifying GrA channel parameters: conductance and lifetime. The inset trace shows GrA current recording, highlighting measured channel parameters: conductance (current amplitude) and lifetime (channel duration). Created with biorender.com. (**B**) Representative current records of the GrA channels reconstituted into a planar lipid bilayer made from lipid mixture of DOPE/DOPG/CL (0.75:0.2:0.05 w/w) before (control) and after addition of 0.5 µM and 1 µM fingolimod to the *cis* side of the membrane. The membrane bathing solution contained 0.1M KCl buffered by 5 mM HEPES at pH 7.4. A voltage of 100 mV was applied. The dashed black line indicates the zero current level; blue dashed lines indicate single channel amplitudes. Addition of fingolimod decreases channel conductance and increases lifetime. For data analysis, the current records were digitally filtered at 5 KHz using a low-pass Bessel (8-pole) filter. (**C**) GrA single-channel conductance normalized versus control conductance at different fingolimod concentrations in 1.0 and 0.1 M KCl. (**D**) GrA lifetime as a function of fingolimod concentration in 1.0 and 0.1 M KCl. Lifetimes were normalized to control values in the absence of fingolimod. Data points represent mean ± SD of 3 independent experiments. Other experimental conditions are as in (B).

### Fingolimod alters planar membrane properties

To investigate the effect of fingolimod on bilayer membrane properties, we measured grA channel conductance and lifetime at increasing concentrations in planar lipid bilayers composed of an *E. coli*-mimicking phospholipid mixture (75% PE, 20% PG, 5% CL), representing the inner membrane and inner leaflet composition of the outer the membrane (Fig. 2B).

Normalized grA channel conductance shows a progressive reduction with increasing fingolimod concentration (0.5–2 µM) in both 0.1M and 1M KCl solutions a salt and dose-dependent manner (Fig. 2C). This reduction is more pronounced at low salt (0.1 M KCl, blue circles), due to greater surface charge neutralization by the positive charges of fingolimod and decrease of the K^+^ concentration near the channel entrance. At high salt (1 M KCl, red circles), this effect is diminished (22). Normalized grA lifetime increases in a dose-dependent manner (Fig. 2D). These results indicate that the membrane hydrophobic core properties are modified upon fingolimod partitioning into the membrane, as shown in the schematic in Fig. 2A, in a manner that stabilizes the grA conducting dimer, resulting in a prolonged lifetime compared to the control.

### Fingolimod permeabilizes *E. coli* and *P. aeruginosa* membranes

Previous work with fingolimod and sphingosine, a structural analog of fingolimod, has demonstrated their membrane permeabilizing activity against *S. aureus* and *E. coli*, respectively (8). Here, we tested propidium iodide (PI) uptake, a membrane-impermeable dye whose fluorescence quantum yield increases markedly upon DNA binding (27). Therefore, bright fluorescence signals indicate membrane permeabilization, as PI gains entry into the cell and is able to bind to genomic DNA. *E. coli* or *P. aeruginosa* grew in the presence of increasing doses of fingolimod (0, 10, 30, 40 µM) for one hour, followed by assays for propidium uptake. As shown in Fig. 3, both *E. coli* and *P. aeruginosa* displayed a dose-dependent increase in PI fluorescence, indicating progressive membrane permeabilization with increasing concentrations of fingolimod. Notably*, E. coli* exhibited slightly higher PI uptake compared to *P. aeruginosa* at the highest (40 µM) tested concentrations, suggesting a subtle species-specific preference for *E. coli*.

**Figure 3.**
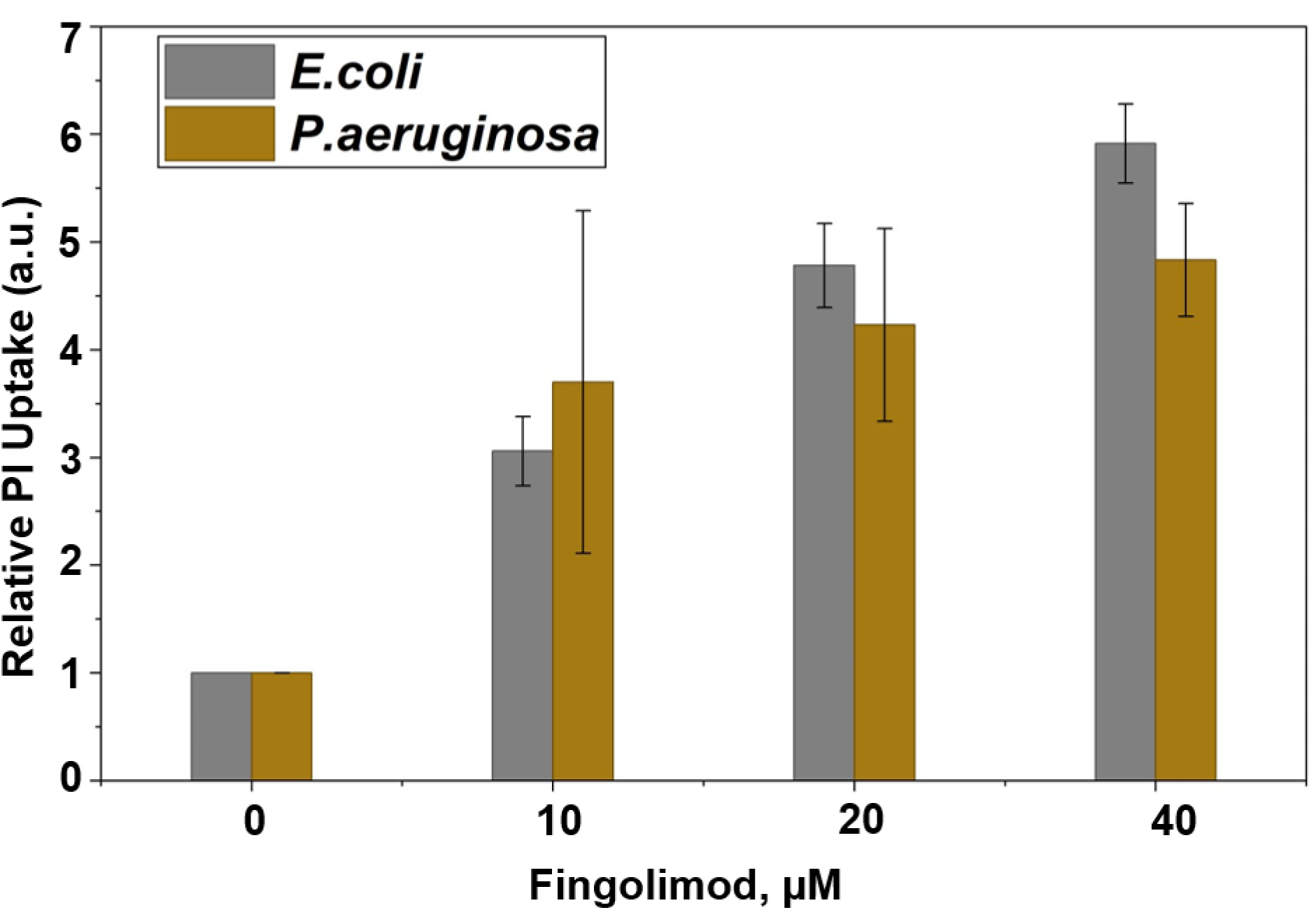
Fingolimod induces membrane permeabilization in *E. coli* and *P. aeruginosa*. *E. coli* and *P. aeruginosa* cultures were treated with increasing concentrations of fingolimod (FL; 0, 10, 20, and 40 µM) for 1 hour at 37 °C, followed by incubation with propidium iodide to assess membrane integrity. PI uptake was quantified fluorometrically as an indicator of membrane permeabilization. A concentration-dependent increase in PI fluorescence was observed in both *E. coli* and *P. aeruginosa*, indicating enhanced membrane disruption. Bars represent mean ± SD from three biological replicates.

### Fingolimod permeabilizes planar lipid membranes by forming lipidic pore-like structures

Our observation that fingolimod displays a dose-dependent response in propidium uptake in *E. coli* and *P. aeruginosa* (Fig. 3) is consistent with a recent observation that fingolimod can disrupt *S. aureus* bacterial membrane integrity (8). These results suggest that the antimicrobial activity of fingolimod is associated with bacterial membrane perturbation, but the exact mechanism remains unknown.

To determine whether fingolimod can directly compromise the barrier function of lipid membranes in the absence of grA or other peptides, we performed experiments using planar lipid bilayers of the same lipid composition as in experiments with grA using micromolar drug concentrations. The representative current traces of the fingolimod-induced transient conductance events, obtained at three different concentrations, under application of 60 mV voltage, are shown in Fig. 4A. At 4 µM of fingolimod (left panel), the current amplitude is mostly low, ∼5-7 pA, suggesting relatively small conductive defects. Increasing the fingolimod concentration to 8 µM (middle panel) resulted in higher current fluctuations up to 200-250 pA, whereas 16 µM fingolimod (right panel) induced a large-scale, up to 750-800 pA, membrane permeabilization, suggestive of a sustained lipidic pore formation.

**Figure 4.**
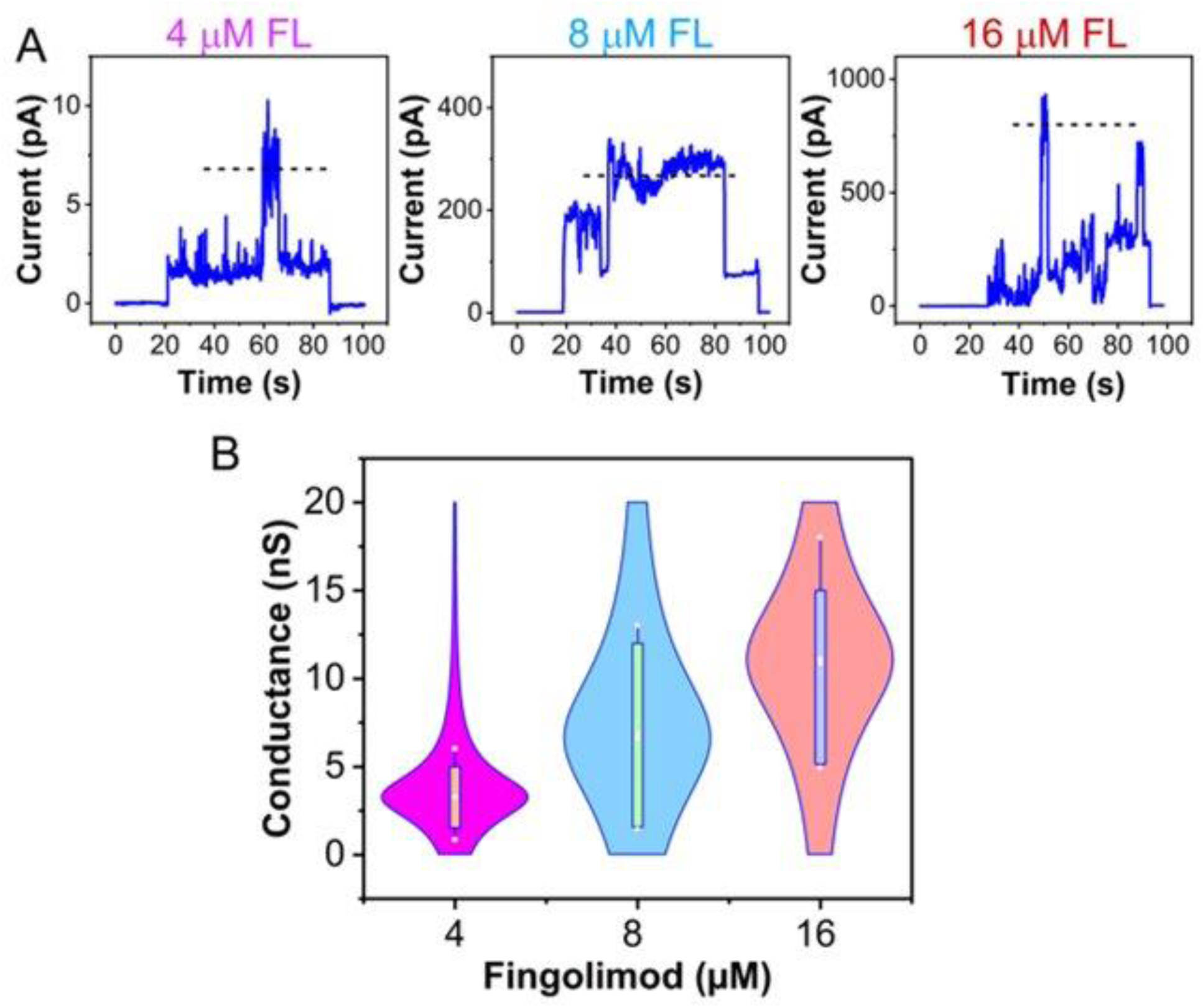
Fingolimod permeabilizes planar lipid membranes at a higher concentration. **(A)** Representative current records obtained on the same PLM at increasing concentrations of fingolimod as indicated, at 60 mV of applied voltage. At 4 µM fingolimod, transient and small-amplitude pore-like events (< 7 pA) were observed, indicative of minor bilayer destabilization. At 8 µM, sustained large-conductance pore formation was evident (∼250 pA), suggesting stable lipidic pore formation. At 16 µM, fingolimod induced high-amplitude, irregular current bursts (∼800 pA), consistent with large-scale membrane permeabilization. Experimental conditions as in Figure 2. **(B)** Violin plot showing the distribution of fingolimod-induced conductance at 4, 8, and 16 µM of fingolimod. The width of each violin reflects the probability density of measured conductance values, with white dots marking the median and boxes indicating interquartile ranges in 6 experiments. A progressive increase in both median and range of conductance is evident with increasing drug concentration.

Notably, fluctuations were rapid, transient, and often reversible at 4 and 8 µM of fingolimod, under the same applied voltage. To show the broad range of membrane conductance at different voltages, we compiled single-event data from multiple traces and summarized them in a violin plot (Fig. 4B), which shows the concentration-dependent conductance heterogeneity. Collectively, these findings suggest the dynamic formation and disappearance of fingolimod-induced pores in the membrane. A question arises whether fingolimod is compromising the lipid bilayer integrity or promoting lipidic pore formation by interacting with the lipid components through an oligomerization-like mechanism (28–30).

### Fingolimod binding to the planar lipid membranes studied by Bilayer Overtone Analysis (BOA)

BOA is a method that allows direct measurements of the binding kinetics of various compounds to planar lipid membranes (31). BOA provides a real-time readout on the transmembrane potential (ΔΨ) by measuring the second harmonic response to the periodic electrical signal, caused by asymmetrical charge distribution between two lipid leaflets (31–34). Fingolimod, which carries a net positive charge due to the presence of a primary amine group, is expected to create an asymmetry in the transmembrane potential when bound to one side of the membrane.

Subsequent additions of fingolimod to the *cis* side of the planar membrane formed from the same lipid mixture used in the experiments with grA, resulted in a stepwise increase in ΔΨ, indicating an asymmetric accumulation of positively charged species on the *cis* side of the membrane (Fig. 5A). Fingolimod’s cationic headgroup can mediate interactions with anionic lipids DOPG and CL, creating an asymmetric charge distribution across the bilayer. Interestingly, the ΔΨ response kept increasing gradually up to 4 µM of fingolimod and sharply declined following the 8 µM addition (Fig. 5A). This behavior indicates a transition from fingolimod binding to (and staying at) the *cis* side of the membrane to its translocation to the opposite, *trans* side bilayer, thus eliminating the charge asymmetry and, consequently, bringing ΔΨ down to zero at > 4 µM of fingolimod. This process was accompanied by a rapid increase in membrane conductance, which was simultaneously measured by BOA (Fig. 5B) on the same membrane. The simultaneous dissipation of transmembrane potential and increase of conductance indicates the translocation of fingolimod to the opposite, *trans* side of the bilayer through the formation of the transient pores and subsequent membrane rupture (indicated by the red arrows in Fig. 5A, B).

**Figure 5.**
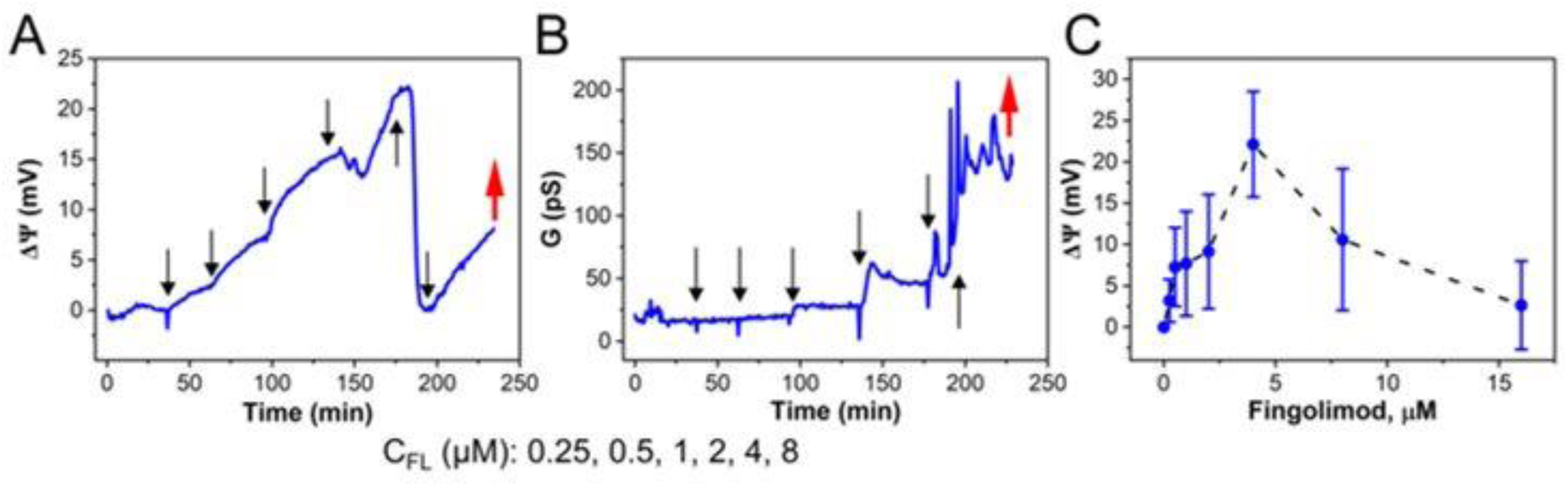
Fingolimod membrane binding measured by BOA suggests its autocatalyzed translocation. (A, **B)** Representative time courses of transmembrane potential (A) and conductance (B) change upon fingolimod addition to the *cis* side of the same planar membrane made from DOPE/DOPG/CL (0.75:0.2:0.05 w/w) in 0.1 M KCl, pH 7.4. Stepwise additions of fingolimod are shown by black arrows; the final concentrations of fingolimod are indicated. Increasing concentrations of fingolimod produced a progressive positive shift in ΔΨ, with a sharp decrease in the transmembrane potential after 4 µM fingolimod addition, followed by an increase in positive potential after 8 µM, and subsequent membrane rupture indicated by the red arrow (A). Relative membrane conductance remained stable at ∼ 15 - 25 pS up to 4 µM fingolimod addition and sharply increased after 8 µM fingolimod addition, indicative of pore formation (B). **(C)** Average ΔΨ as a function of fingolimod concentration. Data points represent the mean ΔΨ values at each fingolimod concentration, with error bars indicating ±SD for 15 independent experiments. A progressive increase ΔΨ is observed with increasing fingolimod concentration up to ∼4 µM, followed by a reverse ΔΨ to a baseline at higher fingolimod concentrations, which suggests fingolimod equilibration between the *cis* and *trans* lipid leaflets of the membrane.

The average binding curve for fingolimod is shown in Fig. 5C, demonstrating the transient character of fingolimod-membrane interaction when the binding regime at low fingolimod concentrations is followed by the translocation regime at the increased fingolimod concentration.

### Fingolimod-induced pore formation is pH dependent

Studies of gram-positive bacteria by Shang et al (8), suggested that fingolimod can compromise the structural integrity of bacterial membranes in a pH-dependent manner. Another recent study has demonstrated that CADs tend to accumulate within acidic cell organelles such as lysosomes, as the low pH environment favors the protonation of their amine groups (35).

In turn, we investigated the impact of fingolimod ionization on its membrane-binding behavior using BOA and conductance measurements at acidic (pH 5.5) and basic (pH 9.5) conditions, considering the reported fingolimod pKa ∼8.0. At pH 5.5, when fingolimod is mostly protonated, subsequent additions of drug (0.25–16 µM) to the planar membrane made from the same lipid mixture, induced a concentration-dependent increase in transmembrane potential, reaching over 10 mV after 2 µM of drug addition (Fig. 6A). The reverse of ΔΨ back to zero, followed by addition of 8 and 16 µM fingolimod (Fig. 6A), was accompanied by a sharp increase in ion conductance (Fig. 6C), consistent with membrane destabilization and lipidic pore formation (Fig. 4 A-B). In contrast, at pH 9.5, fingolimod, being in its neutral, deprotonated form, influenced neither the ΔΨ (Fig. 6B) nor membrane conductance (Fig. 6D) upon drug addition up to 16 µM, revealing the absence of fingolimod-membrane interaction or perturbation.

**Figure 6.**
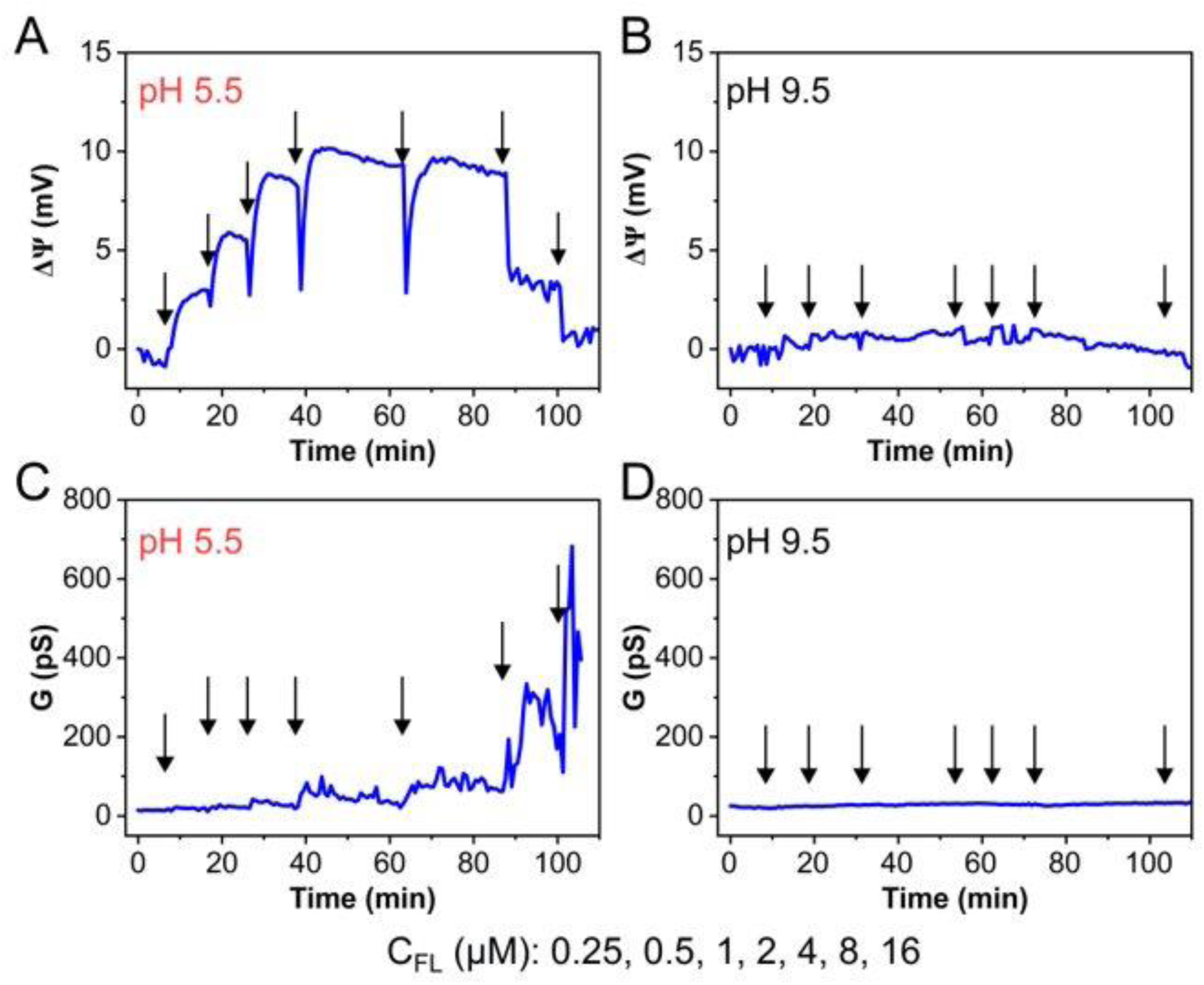
Fingolimod-membrane binding is pH-dependent. Representative time courses of transmembrane potential and conductance changes measured by BOA at different fingolimod concentrations at pH 5.5 (A, C) and pH 9.5 (B, D). **(A, B)** Sequential additions of fingolimod (0.25–16 µM) at pH 5.5 produced progressive shifts in the positive ΔΨ in a dose-dependent manner up to 4 µM, followed by progressive reversal in ΔΨ toward a baseline with further additions of fingolimod (A). At pH 9.5, the addition of fingolimod up to 16 µM did not induce a substantial change in ΔΨ (B). **(C, D)** Relative membrane conductance increased up to 700 pS after the addition of 8 µM fingolimod at pH 5.5 (C) and remained essentially unchanged at pH 9.5 (D). All other experimental conditions are as in Figure 4.

### Fingolimod-membrane interaction studied by molecular dynamics simulations

To clarify the molecular details of fingolimod’s membrane activity, we turned to using molecular dynamics simulations. First, molecular dynamics simulations provided us with the structural features of fingolimod/POPC bilayers (20% fingolimod by mole), as well as the sensitivity of fingolimod to local *curvature*. The coupling of fingolimod to curvature is critical because the pore structure we hypothesize is highly curved, and the curvature of its edge in the center of the membrane is net negative (concave, Fig. 7A) for small pores. The approach is described in more detail in the supplemental material (Fig. S1). Briefly, the curvature preferred by the lipid is determined by where it dynamically enriches on thermally excited membrane undulations. This *spontaneous curvature* explains how it supports the finite-sized pores observed in planar lipid bilayers. Pores are routinely simulated in lipid bilayers (36–38), typically by fixing boundary conditions to prevent collapse. Here, we report the redistribution of fingolimod on a tension-stabilized model pore to determine whether it stabilizes pore expansion or small pores.

**Figure 7.**
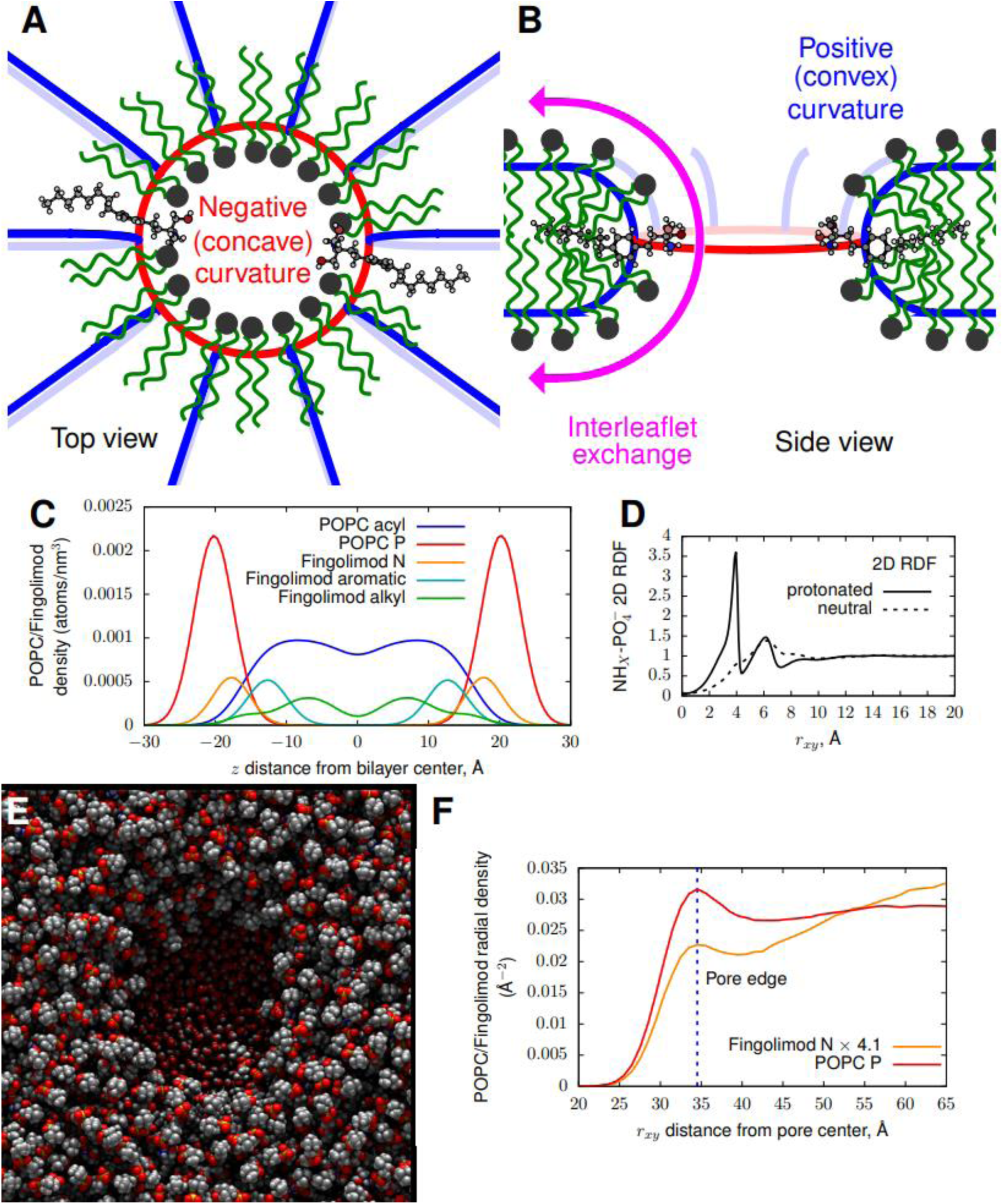
Hypothesized smaller lipidic pore. **(A)** Top view with negative (concave) curvature shown in red, bringing headgroups closer together. **(B)** Sideview of positive (convex) curvature indicated in blue, bringing headgroups apart. Interleaflet continuity facilitating exchange is displayed in purple. Fingolimod molecules, substituting lipids, are shown at the pore wall center where negative curvature is dominating for the small-radius pores. All-atom planar membrane simulations: **(C)** The transverse distribution of the fingolimod charged amine (orange) relative to hydrophobic regions (green/blue) and the POPC phosphate (red); **(D)** 2D (projected) radial distribution functions (RDFs) of the fingolimod amine group and the POPC phosphate group, indicating charged fingolimod has strong interactions with nearby anionic moieties. All-Atom simulation of a large lipidic pore: **(E)** Overhead view of a large, tension-stabilized POPC/fingolimod pore; **(F)** Radial density of fingolimod and POPC, with fingolimod scaled by the ratio of POPC to fingolimod in the simulation. The increased lipid density is at the pore edge due to the inwardly oriented lipids projected into the plane. Fingolimod is depleted at the positively curved pore edge, consistent with its sensitivity to negative curvature.

Second, the simulations demonstrated that model fingolimod strongly favors the hydrophobic bilayer. Fig. 7C shows the location of the fingolimod charged amine group (orange) relative to the POPC phosphate (red). The fingolimod amine is closer to the bilayer center by ca. 2 Å on average, consistent with its analogy to the sphingosine backbone. At no point in the replicas of either the protonated or unprotonated models did fingolimod leave the membrane or flip leaflets in intact bilayers. This finding advocates the importance of the fingolimod-induced transient structural defects (Figs. 4 and 5B) for its inter-leaflet exchange (Fig. 5A).

Fig. 7D shows the radial distribution functions (RDFs) between the fingolimod amine (N atom) and the anionic phosphate moiety of POPC. RDFs indicate correlated positions between two molecular centers, where strong positive correlations indicate a frequently observed interaction and thus strongly suggest stabilization of a complex. The RDF is shown projected into the XY plane for both protonated (solid line) and unprotonated (dashed line) fingolimod models. The protonated form is strongly enriched at an in-plane distance of 4 Å, consistent with a strong electrostatic interaction between the oppositely charged groups. In contrast, the neutral form is moderately depleted (dashed line). Fig. S2 shows the full three-dimensional RDF, meaningful only at short distances, indicating the extreme enrichment of the charge pair.

Third, it was established that fingolimod prefers negative leaflet curvature and is not a pore line-actant. Spontaneous curvature was computed from a simulation using the SPEX method (39, 40) as implemented in the MembraneAnalysis software package (41) (Supplemental material). The spontaneous curvature of both the protonated and unprotonated forms of fingolimod is negative for the model simulated here. Fingolimod interacting with neighboring POPC phosphate groups has spontaneous curvature –0.027 ± 0.002 Å^-1^, with this interacting form present 81% of the simulation time in the protonated model. In a typical phospholipid bilayer, a pore is energetically unfavorable due to the high curvature of bending lipids from one leaflet to another (Fig. 7A, B). Radial expansion of the pore increases the amount of bent lipids linearly, as they line the circumference of the pore. A pore line-actant reduces the *line tension* by stabilizing this interface. While there is substantial negative curvature in small pores (Fig. 7A), the net curvature is positive for large pores, as the convex curvature (Fig. 7B) does not decrease with pore radius. Surfaces can be curved in two directions, such that they have principal curvatures *J*_1_ and *J*_2_ as shown in Fig. 7A, B. Pores, whether fusion/fission pores or single leaflet pores, have both total curvature (*J*_1_ + *J*_2_) that can be positive or negative and Gaussian curvature (*J*_1_ × *J*_2_) that is always negative. Fig. 7E shows an overhead view of a single-bilayer pore created by removing a cylindrical section of a planar membrane and fixing the projected simulation area such that the pore cannot collapse. The leaflet at the pore edge has positive curvature in the radial direction (Fig. 7B), where the leaflet curves from inside to outside. The magnitude of this curvature does not change with pore radius, although the area of the leaflet surface with this curvature increases proportionally to the radius. In the plane of the pore (the red circle in Fig. 7A) curvature is negative. As the pore size decreases, this curvature becomes more negative and can dominate the total local curvature of the lipid in the center of the pore wall. Lipids that prefer positive curvature will enrich at the edges of large pores, consistent with a model of a “line actant” decreasing the effective line tension.

Figure 7F shows the density of fingolimod and POPC in the tension-stabilized pore as a function of distance from the center, with the fingolimod density multiplied by 4.1 so that the average densities are expected to be the same, all else being equal. For this relatively large pore, fingolimod is depleted at the positively curved pore edge, consistent with its preference for negative curvature and contrary to its possible role as a line actant.

## Discussion

In this report, we show that fingolimod, an FDA-approved immunomodulatory drug for multiple sclerosis, exhibits antimicrobial activity by enhancing membrane permeability in *E. coli* and *P. aeruginosa*. To better understand the molecular mechanism of this effect, we used several biophysical approaches to directly probe the interaction between fingolimod and the membrane. Our data support the hypothesis that fingolimod not only forms reversible lipidic pores but also permeates to the opposite membrane leaflet at sufficiently high concentrations. Using grA channels as biosensors in planar lipid bilayers mimicking the phospholipid composition of *E. coli*, we demonstrate that fingolimod partitions into membranes and modulates their properties. We observed dose-dependent changes in grA channel conductance and lifetime. Importantly, fingolimod also permeabilizes unmodified (that is, not containing grA or any other peptide) membranes in a concentration-dependent manner, inducing transient conductance fluctuations of ∼ 3 nS at 4 μM and forming more stable, high-conducting pores of 5-15 nS at higher concentrations (8-16 μM). BOA measurements revealed that binding of fingolimod to one side of the membrane generates a transmembrane potential that decreases upon drug translocation to the opposite membrane leaflet, which most likely occurs through the formation of lipidic pores. Furthermore, we found that fingolimod’s membrane activity is pH-dependent, with protonated fingolimod forming pores, whereas neutral fingolimod shows no measurable membrane activity. These findings are consistent with our hypothesized lipidic pore formation as a molecular mechanism underlying fingolimod’s antimicrobial effects. To our knowledge, this observation is the first report of fingolimod’s ability to form lipidic pores. These results can explain previously reported antimicrobial activities, synergistic effects with conventional antibiotics, and pH-dependent bactericidal properties of fingolimod (7, 8, 13).

All-atom molecular modeling indicated that fingolimod molecules are stable within the bilayer, with no fingolimod molecules leaving the bilayer or flipping leaflets on the microsecond timescale. Radial distribution functions indicate that protonated fingolimod strongly interacts with neighboring anionic moieties, leading to enrichment of this configuration relative to neutral fingolimod. It is the charge-paired interacting configuration of fingolimod that strongly prefers negative curvature, whereas non-interacting fingolimod is enriched at positive curvature. The net-negative curvature preference of fingolimod is consistent with its depletion in large, tension-stabilized bilayer pores (which have net positive curvature). Small-radius pores, in which the concave principal curvature is greatest, have net negative curvature and are thus stabilized by fingolimod. We note that our simulation results do not yield a small stable pore. This is likely due to the timescale for formation, as experiments indicate the pores form slowly compared with the simulation duration (microseconds, compared with longer experimental variations). Constraining a small pore and allowing fingolimod to enrich at this pore may yield its structure, potentially by applying the methodology of (42). Simulation results are consistent with our experimental results, indicating that the fingolimod pores are relatively small. Protonated fingolimod shows strong interactions with neighboring anionic moieties, forming complexes that prefer negative curvature consistent with a small pore. By establishing continuity between leaflets, a pore allows simple diffusion of fingolimod between the outer and inner leaflets, as observed in the experiments on planar lipid membranes upon 4 µM fingolimod addition. Analogous to mechanisms suggested for pore formation by amphipathic antimicrobial peptides in cell membranes (43), we propose a model where transient membrane defects in which lipid headgroups line the pore interior alongside fingolimod molecules lead to lipidic pore formation (Fig. 7A, B). These lipidic pores differ fundamentally from protein channels in that they lack a fixed protein scaffold. Instead, they are dynamic, self-assembled structures stabilized by the curved lipid-water interface.

While our planar lipid bilayer studies provide detailed biophysical characterization of fingolimod-membrane interactions, several questions remain. The lipid composition of the model membranes mimics that of the *E. coli* membrane. However, it does lack the full complexity of bacterial membranes, including proteins, lipopolysaccharides (in gram-negative bacteria), peptidoglycan-associated lipids, and membrane microdomains (44–47). Understanding how these components influence fingolimod pore formation in native bacterial membranes will require future work. Our studies are also limited to symmetric bilayers, in contrast to bacterial membranes that have trans-bilayer lipid asymmetry, which may also affect pore formation (48–51).

Membrane permeabilization by fingolimod may enhance the entry of co-administered antibiotics into bacterial cells, improving their efficacy even against resistant strains. This synergy may be particularly relevant for gram-negative bacteria, where the outer membrane serves as a major permeability barrier limiting antibiotic access. Fingolimod-induced membrane pores could compromise this barrier, sensitizing gram-negative pathogens to antibiotics that would otherwise be excluded. This provides a molecular mechanism linking fingolimod’s chemical structure to its antimicrobial activity. The fingolimod’s amphipathic structure, having a flexible hydrophobic tail similar to a fatty acid and an ionizable cationic headgroup, enables fingolimod to insert into lipid bilayers and stabilize curved lipid-water interfaces, mediating pore formation (Fig. This structure-function relationship provides a starting point for designing fingolimod derivatives with enhanced antimicrobial potency or altered selectivity.

Fingolimod is part of a larger class of cationic amphiphilic drugs (CADs) that includes tricyclic antidepressants, selective serotonin reuptake inhibitors, and other antipsychotic compounds (3, 4). Fingolimod’s ability to perturb membrane structure and form lipidic pores raises the possibility that pore formation may occur with other CADs that have antimicrobial activity. Future studies are needed to determine if other CADs have lipidic pore-forming activity and, therefore, represent a generalizable property of CAD subgroups or a unique feature of fingolimod. Understanding the structure-activity relationships that govern membrane interactions and pore-forming activity could inform rational drug design strategies. For human-targeted therapeutics, this knowledge could aid in modifications that retain desired therapeutic activity while minimizing unwanted effects on bacterial membranes. Conversely, for antimicrobial drug development, the pore-forming mechanism could be exploited by designing new membrane-targeting antibiotics based on the fingolimod structure or by using pore-forming molecules to enhance membrane crossing of other antimicrobial agents. Taken together, our results uncover a previously unknown antimicrobial mechanism for fingolimod involving direct membrane permeabilization via lipidic pore formation.

## Material and Methods

### Bacterial Growth Assays

Overnight cultures of *E. coli* and *P. aeruginosa* were diluted to OD_600_ = 0.1 in LB medium. For the drug screening, *E. coli* was treated with 40 µM of each 25 CADs in 5 mL cultures. For dose-response assays, cultures were treated with 0, 10, and, 40 µM fingolimod. Cultures were plated in 96-well plates (200 µL per well), and OD_600_ using a Tecan Infinite 200 Oro Plate reader as described previously (5).

### Gramicidin A measurements

1,2-dioleoyl-sn-glycero-3-phosphoglycerol (DOPG), dioleoyl-phosphatidylethanolamine (DOPE) and cardiolipin (CL) were purchased from Avanti Polar Lipids, Inc. Alabaster, AL. Gramicidin A (grA) was generously gifted from O. S. Andersen, Cornell University Medical College. Planar lipid bilayers were formed using DOPE/DOPG/CL (0.75:0.2:0.05 w/w) from the opposition two monolayers across a ∼125μm aperture in a 15-μm-thick Teflon partition separating two ∼1.5-mL compartments as described elsewhere (52). Membrane bathing solutions contained 0.1M KCl or 1M KCl buffered with 5 mM HEPES at pH 7.4. GrA dissolved in ethanol solutions was added to both compartments of the chamber. Fingolimod (FL) dissolved in DMSO was added to the *cis* side of the membrane. Single channel measurements were acquired using an Axopatch 200B amplifier (Axon Instruments, Inc., Foster City, CA, USA) in the voltage clamp mode. Sampling was done at 200 Hz frequency with a low-pass Bessel filter at 0.05 kHz. Later, the records were digitally filtered at 0.1 kHz using the Bessel-8 pole filter using the Clampfit 10.7 software as previously described (52). GrA channel lifetimes and mean conductance were analyzed with the previously reported protocol (52, 53).

### Bilayer Overtone Analysis (BOA) Measurements

BOA measurements were conducted as described earlier (31) using a Stanford Research Systems 830 lock-in amplifier. The trans membrane potential Ψ reports the asymmetry between two leaflets of the lipid bilayer. In our experiment, Ψ was quantified by analyzing the second harmonic wave of bilayer current in response to an application of sinusoidal voltage across the lipid bilayer (34). Data are calculated as ΔΨ = Ψ−Ψ_t=0_, where positive value of ΔΨ is analogous to the increase of positive charge on the *cis* side of the membrane. Membrane capacitance and ΔΨ were recorded at one-minute intervals using custom-built software written in Python 3.11 (31). ΔΨ values were averaged over 10 subsequent measurements following a steady-state level, operationally defined by ΔΨ variations within ±0.5 mV.

### Propidium Iodide (PI) Uptake Assay for Bacterial Membrane Permeabilization

Overnight cultures of *E. coli* and *P. aeruginosa* were diluted to an OD_600_ of 0.2 in fresh LB medium and grown for 2 hours at 37°C. Cells were then collected at OD_600_ ∼0.3 and incubated with increasing concentrations of fingolimod (0, 10, 20, 40 µM) for 1 hour at 37°C. Following treatment, cells were washed three times with 10 mM HEPES buffer (pH 7.4) to remove residual drug. For each sample, 100 µL of washed cells were mixed with 100 µL of HEPES buffer containing 2.5 µM propidium iodide (PI), a membrane-impermeant DNA-binding dye. Samples were transferred to black-walled 96-well microtiter plates, and fluorescence was measured using a Tecan Infinite 200 Pro plate reader (excitation/emission 535/617 nm). PI uptake was calculated as: (F_sample-PI_) – (F_sample-PI_)/ (F_buffer+PI_)-(F_buffer-PI_), where *F* refers to fluorescence intensity.

### Membrane conductance measurements in the absence of grA

Planar lipid membranes (PLM) were formed as described for grA and BOA measurements. Data were filtered by a low-pass 8-pole Butterworth filter (Model 900 Frequency Active Filter, Frequency Devices) at 15 kHz, digitized with a sampling frequency of 50 kHz, and analyzed using pClamp 10.7 software (Axon Instrument). For data analysis, digital filtering using a 0.5 kHz low-pass Bessel filter was applied. After membrane was formed, recordings were taken in the absence of fingolimod for ∼30-40 minutes, then fingolimod was added to the *cis* side. A voltage in the range of 30-100 mV was applied to observe membrane permeabilization.

### Molecular dynamics simulations

#### Planar membrane simulations

All simulations were performed at 30C using the Amber (22/PMEMD) simulation package (54) run on a single GPU. Simulations were built on the CHARMM-GUI web platform, including the parameterization of fingolimod (using the CHARMM-GUI Ligand Reader & Modeler module using the CGenFF forcefield). Initial relaxation used the standard CHARMM-GUI “equilibration” steps, while dynamics used the recommended parameters consistent with the forcefield and which are included as input files from CHARMM-GUI (version 3.7). Four replicas of both the protonated and unprotonated fingolimod were built and simulated for at least 4 microseconds. An additional pure POPC bilayer (200 lipids/leaflet) was simulated for one microsecond to compare the area-per-lipid with and without fingolimod (see supplemental material). The neutral surface atom and curvature sensitivity were computed using the MembraneAnalysis Julia package (41). Interacting fingolimod/POPC pairs were identified by first computing the 3D fingolimod-N/POPC-P RDF (Fig. S2), then choosing an upper limit (5 Å) that captured the highly enriched first peak.

#### Pore simulations

For the pore simulations, a patch of the planar protonated fingolimod/POPC bilayer was removed following at least 50 nanoseconds of simulation from each planar replica. With the box area fixed, the pore was unlikely to collapse and so persisted through the simulation, allowing the enrichment or depletion of fingolimod at the pore edge to be quantified. Nevertheless, a pore in one replica collapsed, yielding three metastable pores with lifetimes up to three microseconds. To quantify the distribution of fingolimod in the pore, the bilayer was first centered to place the pore at the origin, then fingolimod and POPC were counted in a histogram starting from the origin, using the neutral surface atom as quantified by Membrane Analysis for the planar simulations above.

## Acknowledgments

Research is supported by grants from the National Institutes of Health National Cancer Institute (R15CA267890-01), the National Science Foundation (CCF PIPP Grant 2200138), and The Catholic University of America Innovation Seed Fund to J.S.C., and by the Intramural Research Program of the *Eunice Kennedy Shriver* National Institute of Child Health and Human Development, National Institutes of Health (grants ZIA HD000072 to S.M.B. and 1ZIA-HD008955 to A.S.). Molecular dynamics simulations were performed using the computational resources of the NIH High-Performance Computing Biowulf cluster (https://hpc.nih.gov). The contributions of the NIH authors are considered Works of the United States Government. The findings and conclusions presented in this paper are those of the authors and do not necessarily reflect the views of the National Institutes of Health or the U.S. Department of Health and Human Services.

## Supplemental Figures and Table

**Table S1.**
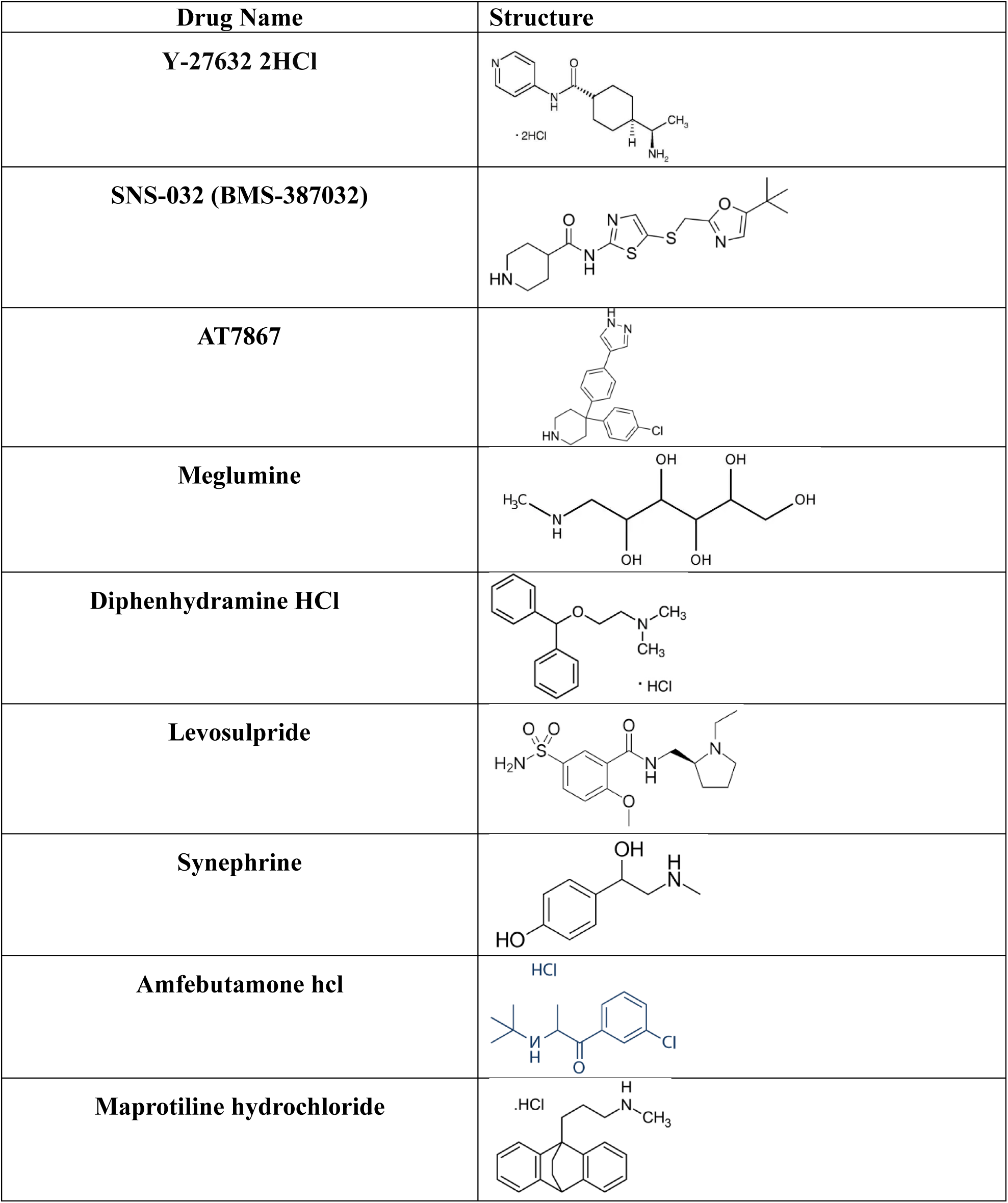
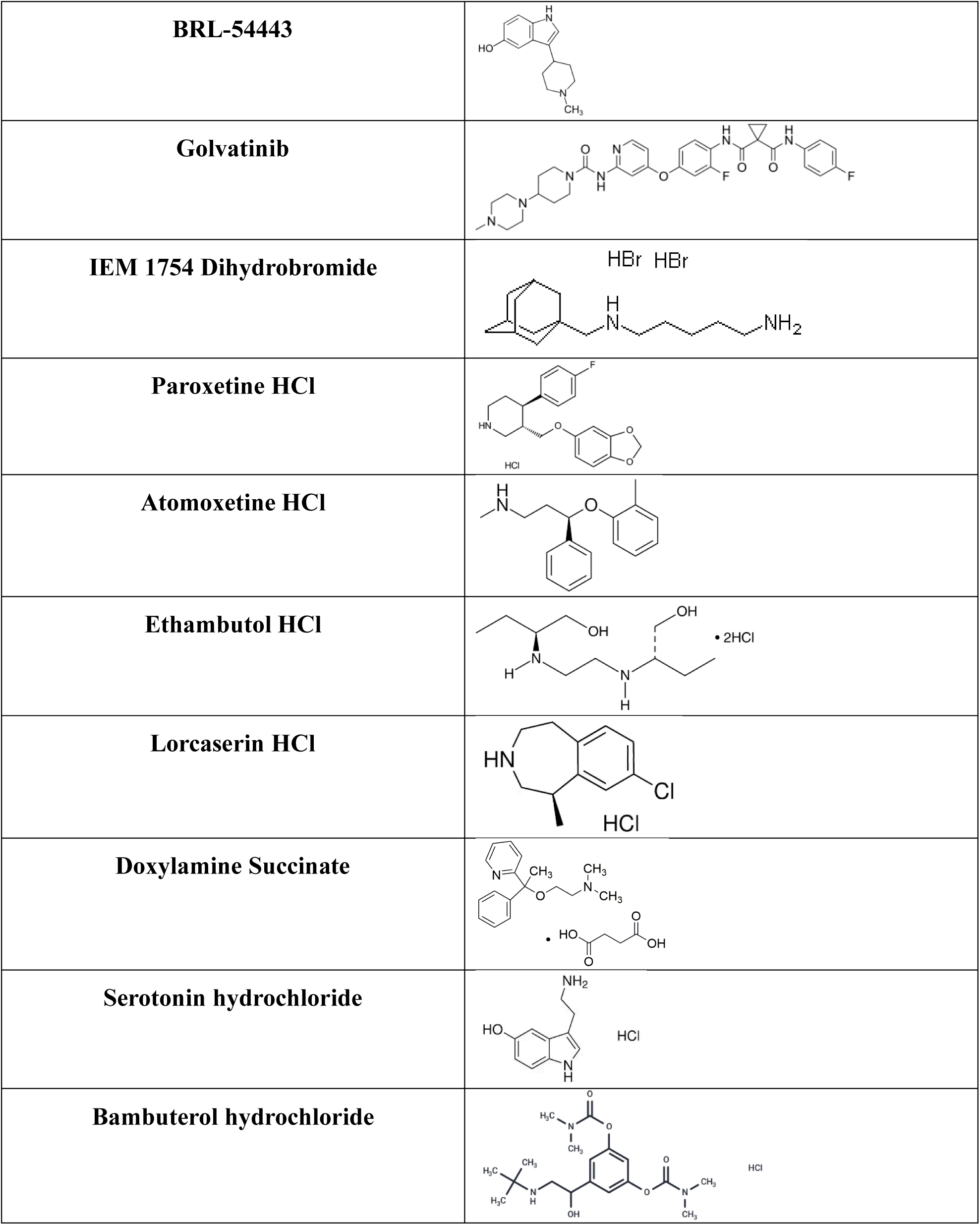
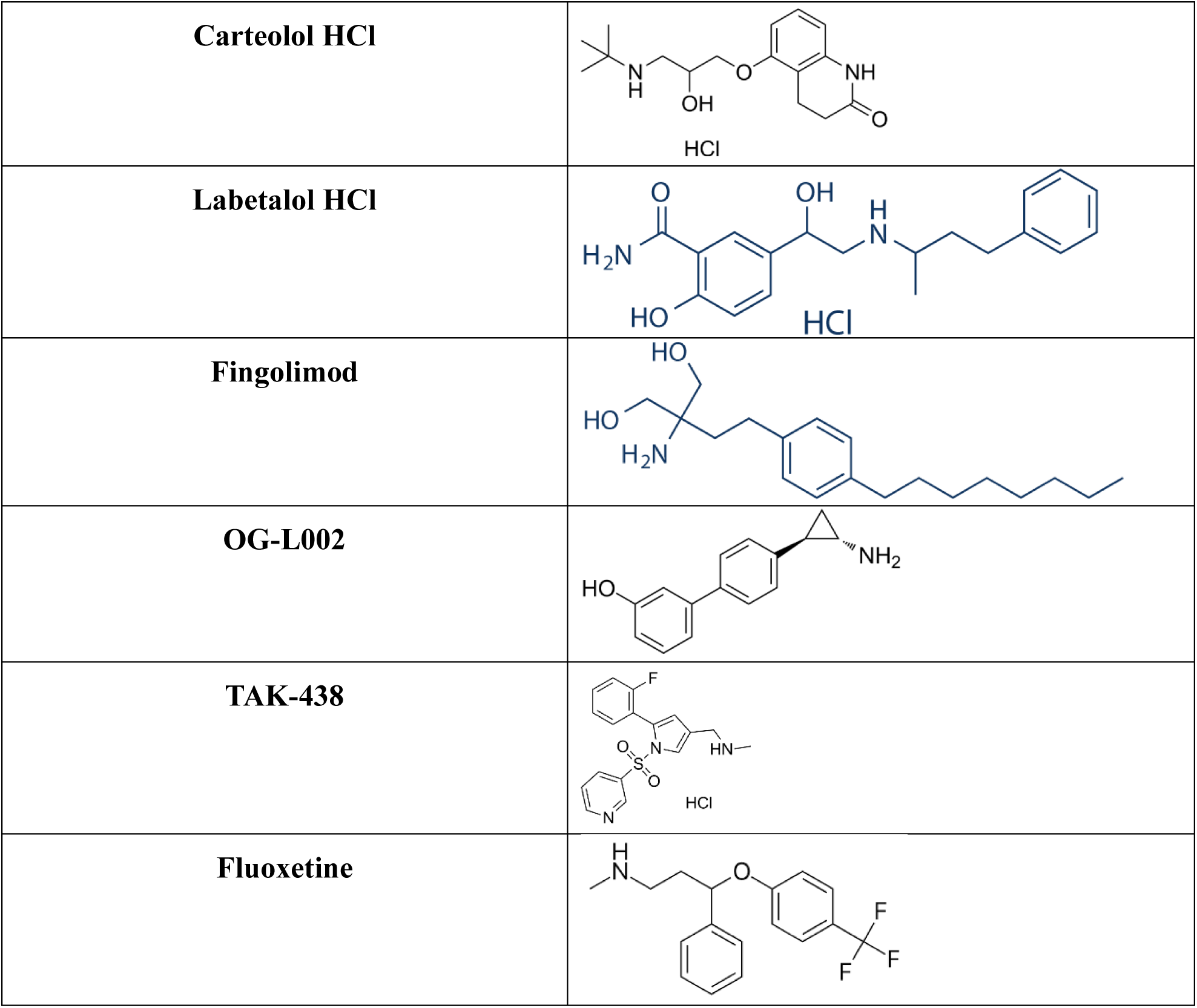
Screening of 25 CAD compounds and their structure.

## Approach to extract spontaneous curvature

**Fig. S1.**
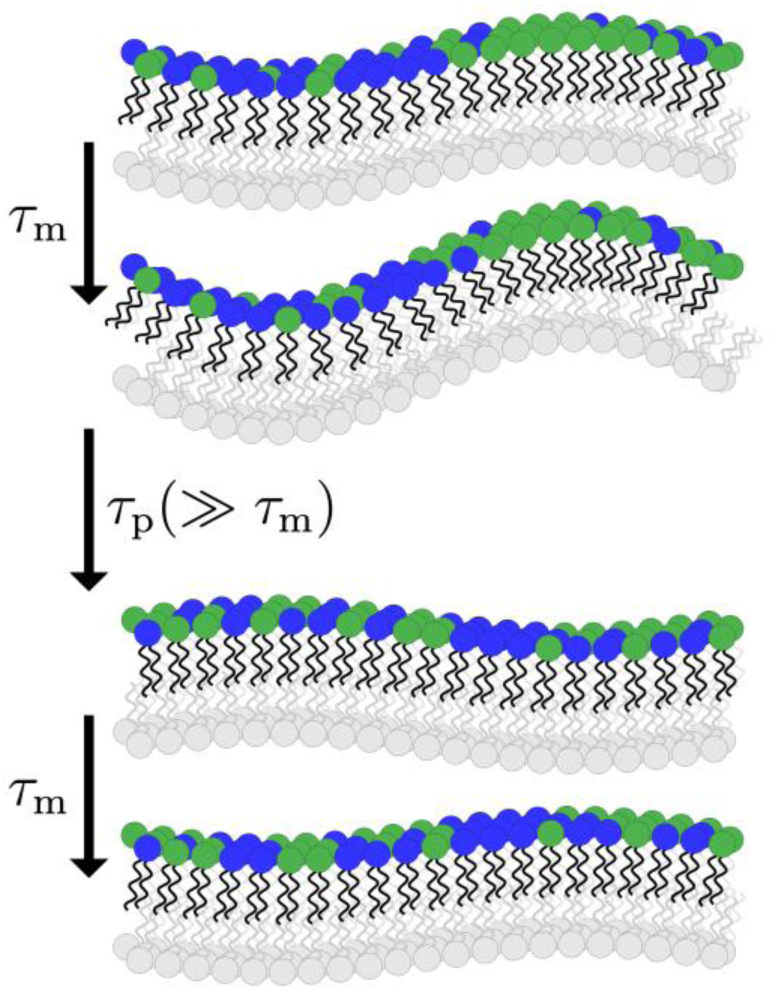
The green lipid has *positive* (convex) spontaneous curvature (*J*_0_) and so prefers the hills of a membrane undulation. Averaging over membrane undulations (with relaxation time *τ*_m_) and lateral redistribution of the lipids (with relaxation time *τ*_p_), the curvature sampled by the green lipid on a single undulating mode, 〈*J*〉, will be proportional to its spontaneous curvature 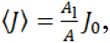, where *A*_1_ is the area-per-lipid of the green lipid and the *A* is the total bilayer area. For a given wavenumber 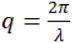 (*λ* is the undulation wavelength), the product *A*〈*J*〉 is invariant to system size, assuming the lipid has a point-like effect on the bilayer. Adapted from Ref. SA, Figure 1.

Fig. S1 illustrates the framework used to extract lipid curvature preference (the spontaneous curvature). As an example, the green lipid prefers positive curvature, the convex hills of the membrane undulation. A random inhomogeneous distribution (shown on the first line) of the green lipid induces an undulation that favors that lipid, shown on the second line. Over time, lateral distribution relaxes and the undulation decays. Both the undulation and inhomogeneous lateral distribution are weaky, thermally excited distributions. Yet the stochastic distributions are not independent, they are *coupled* by the green lipid’s spontaneous curvature. At the scale of these simulations, the bilayer relaxes much more quickly (10s nanoseconds) compared to the lateral compositional variation (approximately a microsecond).

The experienced curvature, 〈*J*〉, is then evaluated by averaging over each fingolimod molecule at each frame of the simulation, taking into account its lateral position and the curvature at that point, given a Fourier decomposition of the membrane undulations. The equilibrium average will be 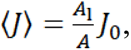 where here *A*_1_ is the projected area (“footprint”) of the lipid or protein. The fraction 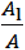 is typically much smaller than 1, for example, in a simulation with 400 lipids per leaflet, the fraction will be 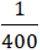 if all lipids have the same *A*_1_. Here the simpler fingolimod with a sphingosine backbone and one acyl chain is smaller than the zwitterionic, double-tailed POPC.

## Radial distribution function (3D)

**Fig. S2.**
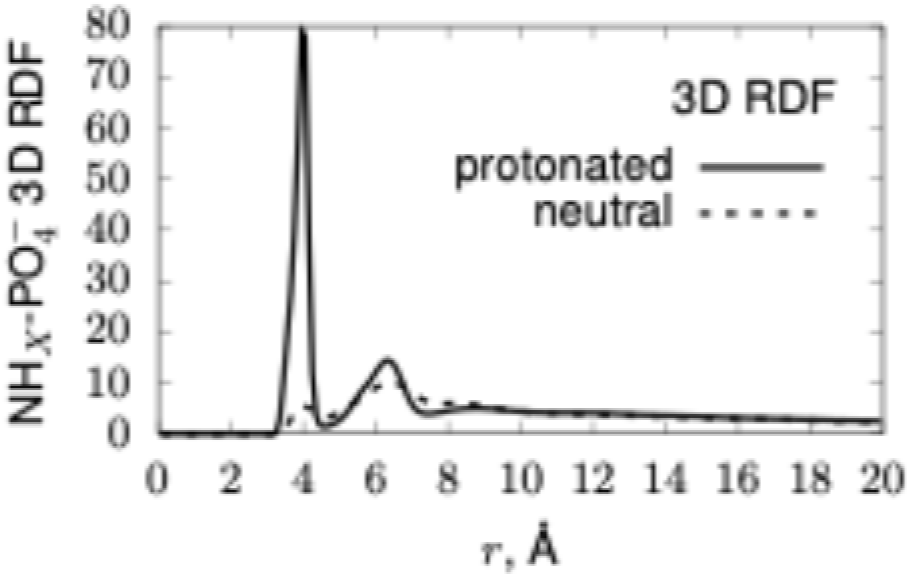
Fingolimod (amine) to POPC (phosphate) radial distribution function (here in 3D) of both the protonated (solid) and unprotonated (dashed) forms.

The radial distribution function (RDF) is shown in Fig. S2. The peak at approximately 4 Å indicates the preferred separation between the positively charged amine group and the negatively charged phosphate (considering the N and P atoms, respectively). The normalization of this RDF assumes a uniform (three-dimensional) distribution of atoms, whereas the structure imposed by the bilayer is pseudo two-dimensional. Therefore, the absolute value of the RDF depends on the surface-to-volume ratio and is not mechanistically meaningful. Comparing the neutral and protonated peaks does offer insight into the strength of the interaction for protonated fingolimod. Additionally, the peak and its shape determine the cutoff (5 Å) used to identify interacting pairs of fingolimod and POPC whose curvature sensitivity is quantified in analysis in Fig. S3.

**Fig. S3.**
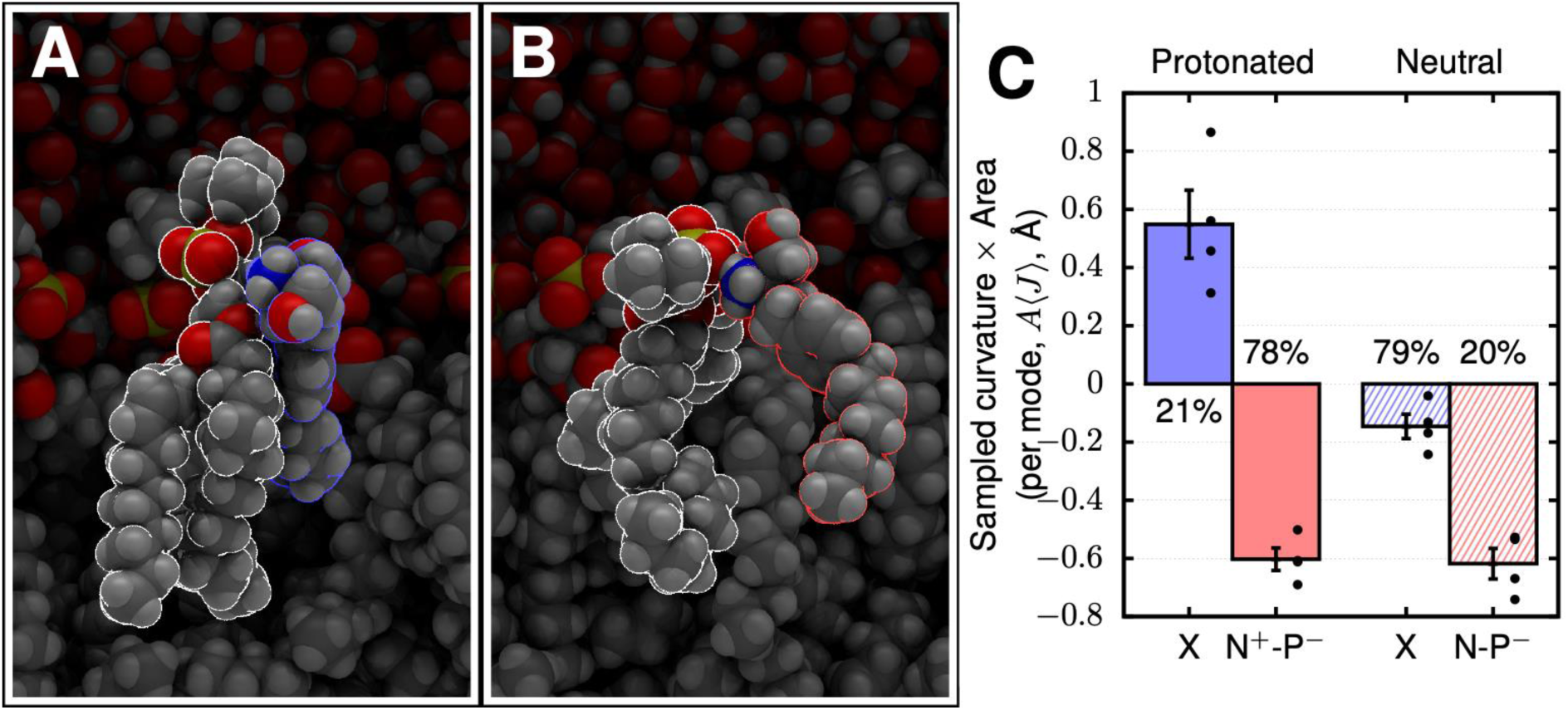
Molecular illustration of proximal but not directly interacting (panel A) and closely interacting (panel B) complexes of protonated fingolimod and POPC. Panel (C) shows the area-weighted curvature *J* sampled by the two forms. The interacting form is the dominant (81%) form of protonated fingolimod (shown in both panels A and B) while the non-interacting form is the dominant species of neutral fingolimod. Error bars report a single standard error.

Fig. S3 shows molecular configurations for non-interacting (panel A, blue) and interacting (panel A, red) fingolimod-POPC pairs. Panel C shows the area-weighted curvature sampled by the configurations. Spontaneous curvature is equal to 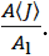 The projected lipid area of fingolimod is difficult to characterize when present in a POPC mixture, as it can easily have a slight condensing effect. For example, the area per lipid of pure POPC we have simulated is approximately 65.3 Å^2^. Subtracting the area of the ca. 300 POPC lipids-per-leaflet in our simulation from the total area, and dividing by the number of fingolimod (∼72), yields a fingolimod area of ∼6 Å^2^, a value that is likely too small due to the condensing effect. Compensating for the condensing effect by taking into account the thickness strain induced by fingolimod (Fig. S4) yields a condensed POPC area-per-lipid of 61.1 Å^2^, implying an area-per-lipid of 22.2 Å^2^, in line with its single acyl chain and small headgroup. With *A*_l_ taken as 22.2 Å^2^, and *A*〈*J*〉equal to -0.6 Å (Fig. S3, panel C), the spontaneous curvature *J*_0_ is -0.027 ± 0.002 Å^-1^.

**Fig. S4.**
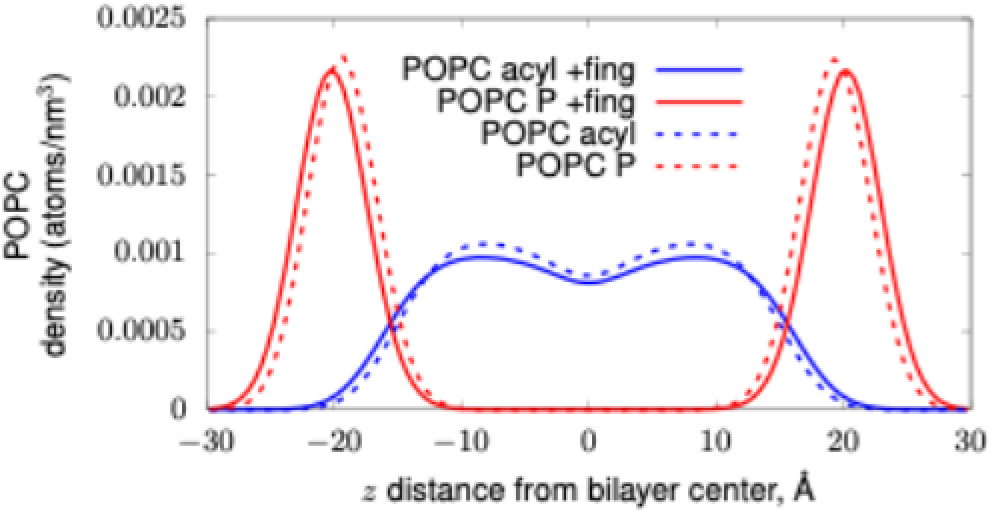
Comparison of the density of phosphate and acyl chain groups with (+fing) and without fingolimod. Modeled protonated fingolimod induced a ∼6% strain in the thickness of the bilayer.

